# The process that the evolution from C_3_ towards C_4_ photosynthesis is driven

**DOI:** 10.1101/2024.04.26.591276

**Authors:** Yong-Yao Zhao, Yao Xin Zhang, Xin Qi Yu, Xin-Guang Zhu

## Abstract

Compared to the plants with C_3_ photosynthesis, the plants with C_4_ photosynthesis have higher light, water, and nitrogen use efficiencies. Though only 3% of higher plants have C_4_ photosynthesis, C_4_ photosynthesis contribute about 23% of terrestrial gross primary productivity. Currently, it remains elusive how the procedure of the evolution from C_3_ towards C_4_ photosynthesis is driven. Here, *Flaveria* genus, the chief model for C_4_ evolutionary researches, was used to study this driving through a combined analysis of ecological, physiological and biochemical data. From proto-Kranz to C_4_-like, results showed that precipitation was negatively associated with the activity of C_4_ metabolic core enzyme and C_4_ flux, respectively. With increased vein density, an increased temperature and precipitation were observed during the earlier evolutionary stage. Our study indicated that decreasing precipitation and increasing temperature accompanied the transition from C_3_ to C_2_ photosynthesis and further transition from C_2_ to developing C_4_ photosynthesis. There was a positive relationship between CO_2_ compensation point, stomatal density and precipitation, supporting the notion that decreasing precipitation enhances photorespiration through reducing stomatal conductance during the evolution. As the driving force from proto-Kranz to C_4_-like, our study highlights that drought dominated rather than heat during C_4_ evolution. This study suggests that the driving force between certain stages during the evolution towards C_4_ photosynthesis are different and increased water availability should be needed in the initial phase.

## Introduction

Photosynthesis undergone evolution, which results in different photosynthetic type, and C_4_ photosynthesis is one of these photosynthetic types (Sage 2004; Langdale 2011). Compared to C_3_ relative, the plant with C_4_ photosynthesis has higher water use efficiency, nitrogen use efficiency and light use efficiency (Vogan and Sage 2011; Taylor et al. 2010). Although only accompanying 3% plant species, C_4_ photosynthesis contributes above 20% of terrestrial gross primary productivity (Still et al. 2003; Sage et al. 2012). The evolution from C_3_ to C_4_ photosynthesis go through multiple stages (C_4_ evolution) (Sage et al. 2012; Sage 2004; Schluter and Weber 2016; Brautigam and Gowik 2016). The first stage of C_4_ evolution is preconditioning in which C_3_ plants acquire new cis-elements (Akyildiz et al. 2007).For this stage, duplication of gene occurs (Akyildiz et al. 2007; Monson 2003), vein density increases (Marshall et al. 2007; Mckown and dengler 2007) and the number of plastids in the bundle sheath increases (Sage et al. 2013). The second stage is proto-Kranz. In this stage, plants still perform C_3_ photosynthesis (Sage et al. 2013) while the number of plastids continue to increase (Stata et al. 2019; Sage et al. 2012) and chloroplast and mitochondria localize to the centripetal wall of bundle sheath cells. The third stage is C_2_ photosynthesis (Type I) (Schulze et al. 2013; Sage et al. 2012; Edwards and KU 1987), which is characterize as the change in the location of GLYCINE DECARBOXYLASE P SUBUNIT (GDCP) and photosynthetic apparatus from mesophyll cell to bundle sheath cell (Schulze et al. 2013; Stata et al. 2019; Sage et al. 2013). The released CO_2_ from photorespiration can be moderately concentrated in the bundle sheath cell (Photorespiratory CO_2_ pump) (Keerberg et al. 2014; Sage et al. 2012). The fourth stage is to induce and develop C_4_ photosynthesis (Type II) (Schluter and Weber 2016; Sage et al. 2018; Nakamura et al. 2013; Mallmann et al. 2014). The establishment of C_2_ photosynthesis could cause the imbalance of ammonia between mesophyll cell and bundle sheath cell, and the induction of C_4_ metabolism is proposed as a mechanism to resolve this trouble (Mallmann et al. 2014). The fifth stage is functionally developed C_4_ photosynthesis (C_4_-like) (Stata et al. 2019). For this stage, though there are still some Rubisco in mesophyll (Taniguchi et al. 2021; Monson et al. 1987; Stata et al. 2019; Sage et al. 2018), various types of indicators at this stage are close to plants with ultimate C_4_ photosynthesis (Vogan and Sage 2011; Ku et al. 1991; Adachi et al. 2023). The final stage is the mature C_4_ photosynthesis (Mature C_4_) (Sage et al. 2012), and mesophyll cell contain few RuBisCO (Stata et al. 2019). These stages in the evolutionary transition between C_3_ and C_4_ photosynthesis are called intermediate stage, and the plants in these stages are intermediate plants (intermediate species).

The evolution from C_3_ towards C_4_ photosynthesis is considered to be driven by enhancing photorespiration, which is promoted by low concentration of atmosphere CO_2_, drought, salt and heat (Sage et al. 2012; Brautigam and Gowik 2016; Sage 2004; Edwards and Smith 2010; Bromham and Bennett 2014). Low concentration of atmosphere CO_2_, drought and salt could decrease the ratio of CO_2_ to O_2_ concentration, thereby enhancing photorespiration (Sage et al. 2012). Increasing temperature enhances photorespiration by reducing the concentration of CO_2_ relative to O_2_ in the chloroplast stroma (Ogren 1984) and decreases the specificity of RuBisCO for CO_2_ relative to O_2_ (Ogren 1984; Sage et al. 2012). Studies have shown that C_4_ evolution are correlated with the decrease in the concentration of atmosphere CO_2_ and the appearance of dry, salty and hot environment (Christin et al. 2008; Edwards and Smith 2010; Edwards and Still 2008; Bromham and Bennett 2014), which suggests they are the driving force for C_4_ evolution. However, all these studies assume that the entire process of C_4_ evolution is under a single driver. Given C_4_ evolution includes multiple phases and the evolution is complicate, it is possible that these phases of C_4_ evolution might be under different driving force rather than the same driving force persistently along the evolutionary road towards C_4_ photosynthesis (Heckmann et al. 2013; Sage et al. 2012).

Plants obtain fitness in each step of C_4_ evolution (Heckmann et al. 2013; Sage et al. 2012). The increased vein density is an important characteristics in the initial stage of C_4_ evolution (Sage et al. 2012; Sedelnikova et al. 2018; Kumpers et al. 2017; Sage et al. 2014) (Mckown and dengler 2007; Marshall et al. 2007; Williams et al. 2013), which might help supply water to the mesophyll and then maintain stomatal opening, since evaporation might be enhanced at this time (Brautigam and Gowik 2016; Sage et al. 2018) (Sage et al. 2012). Therefore, available water content in the environment for this stage may be increased to ensure the fitness of plants (Brautigam and Gowik 2016). However, this contradicts with the notion that the origin of C_4_ photosynthesis is promoted by dry environment (Sage et al. 2018; Edwards and Smith 2010; Sedelnikova et al. 2018). Since C_2_ photosynthesis and C_4_ photosynthesis could largely reduce photorespiration (Sage 2004; Schluter and Weber 2016; Sage et al. 2012), the formation of C_2_ from C_3_ photosynthesis and the transition from C_2_ to developing C_4_ photosynthesis may be driven by the conditions which could enhance photorespiration (Schluter and Weber 2016; Sage et al. 2018; Sage et al. 2012),but the driving forces during these transitions also remain undefined.

*Flaveria* genus is a widely used system to study C_4_ evolution (Sage et al. 2012; Sage 2017; Schluter and Weber 2016; Mallmann et al. 2014; Sage et al. 2013; Vogan and Sage 2011; Nakamura et al. 2013), and the studies on *Flaveria* genus has laid the foundation for the pyramid model of C_4_ evolution (Sage et al. 2013; Vogan and Sage 2011; Sage et al. 2012). The photosynthetic types of *Flaveria* species span entire evolutionary stage towards C_4_ photosynthesis(Sage et al. 2012; Sage et al. 2014; Schluter and Weber 2016; R.K. Monson 1986). *Flaveria* is also the youngest genus among the known C_4_ linage and it’s species has the close relationship (Schulze et al. 2013). These make *Flaveria* genus an ideal model to study C_4_ evolution (Christin et al. 2011; Sage et al. 2012). The ecology for the habitats of intermediate species and C_4_ species could reflect the environmental conditions that drive C_4_ evolution (Sage et al. 2012; Sage et al. 2018), therefore, the ecology of *Flaveria* habitats enable us to finely study the driving force in the evolutionary road towards C_4_ photosynthesis. In this study, we collected the ecological, biochemical and physiological data for different species of *Flaveria* genus to explore the procedure of driving for the evolutionary trajectory towards C_4_ photosynthesis and how these different evolutionary phases help plants gain fitness.

## Results

### 1. Most of *Flaveria* species were distributed inland and a few species were distributed along the coast

The habitats of *Flaveria* species were mapped to the North and South America continents with the longitude and latitude of each *Flaveria* samples downloaded from Global Biodiversity Information Facility (GBIF) (Fig 1A, B). Species were ordered based on the amount of C_4_ flux (Fig 1B, C, supplementary fig 1). We found most *Flaveria* species were located inland (Fig 1A, B). A few species (*F*.*bro, F*.*flo, F*.*lin*) were distributed along the coast (Fig 1B), which was consistent with the previous reports (Powell 1978). During evolution, intermediate species gradually spread to locations with higher latitude (Fig 1B). Compared with the intermediate species, the distribution of C_4_ species was more widespread (Fig 1A, B, C), indicating that adaptive capacity of C_4_ plants is enhanced (Fig1).

**Fig 1.**
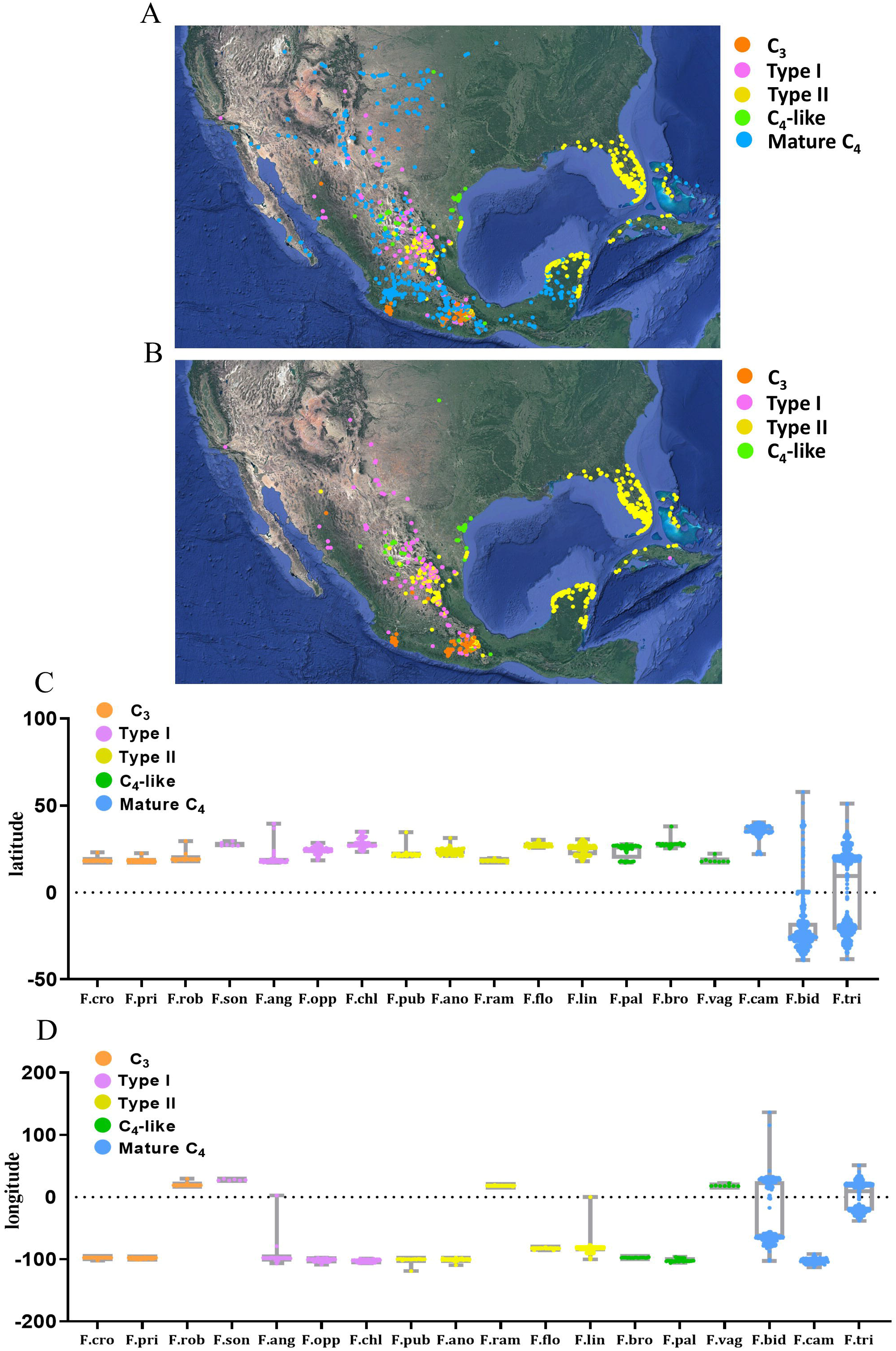
Distribution of the *Flaveria* species (A) The location of *Flaveria* species was mapped to the world map. The main habitat of *Flaveria* is North and South America, which was displayed. (B) The location of *Flaveria* species except C_4_ species was mapped to the world map. (C) and (D) The latitude and longitude of different *Flaveria* species. The latitude and longitude were acquired from GBIF. Different colors indicate different photosynthetic types. *F*.*rob* (n=39), *F*.*cro* (n=48), *F. pri* (n=140), *F*.*son* (n=6), *F*.*ang* (n=50), *F*.*ano* (n=145), *F*.*opp* (n=135), *F*.*pub* (n=25), *F*.*ram* (n=65), *F*.*chl* (n=96), *F*.*flo* (n=56), *F*.*lin* (n=799), *F*.*cam* (n=95), *F*.*bro* (n=19), *F*.*bid* (n=525), *F*.*pal* (n=44), *F*.*tri* (n=1652), *F*.*vag* (n=8).

### 2. The changed pattern for the temperature and precipitation of *Flaveria* growth habitats

The temperature and precipitation of the habitats for each *Flaveria* species were obtained from the worldclim database (Fick and Hijmans 2017). The results from physiological, anatomic, molecular and computational modeling experiments indicate the direction of C_4_ evolution could be described by species’s order towards C_4_ photosynthesis in phylogenetic tree (Figure 2A) (Mckown and dengler 2007; Nakamura et al. 2013; Gowik et al. 2011; Sage et al. 2012; Sage et al. 2018; Schluter and Weber 2016). Along the evolutionary pathway towards C_4_ photosynthesis (Ordinal species towards C_4_ photosynthesis of the phylogenetic tree in Figure 2A), the annual mean temperature, mean temperature of warmest quarter, mean temperature of wettest quarter and max temperature of warmest month first increased and then decreased, though difference in some species habitats is not significant (Fig 2A, supplementary fig 2, supplementary fig 3, supplementary fig 4, supplementary file). Mean temperature of warmest quarter, mean temperature of wettest quarter and max temperature of warmest month increased significantly in the habitat of *F. rob* (C_3_) and *F. son* (Type I) (Fig 2A, B, supplementary fig 2, supplementary fig 3, supplementary file), compared with those of *F. pri* (C_3_). However, these environmental conditions significantly decreased in the habitat of *F. ang* (Type I), compared with *F. son* (Type I) (Fig 2A, supplementary fig2, supplementary fig3, supplementary file). For the mean temperature of wettest quarter, mean temperature of warmest quarter and max temperature of warmest month, compared with *F. son* (Type I), the habitat of species with C_2_ photosynthesis (*F. ang, F. chl, F. pub, F. opp, F. ano, F. ram*) had a lower mean temperature, but the results of statistic test for some species are not significant or in margin (Fig 2A, supplementary fig 3D, supplementary file). The habitats of species *F. flo* (Type II) and *F. lin* (Type II) had significantly higher mean temperature compared to most of other intermediate species (Fig 2A, supplementary file). Max temperature of warmest month for *F. son* (Type II) habitats was extremely high (Fig 2d, supplementary file).

**Fig 2.**
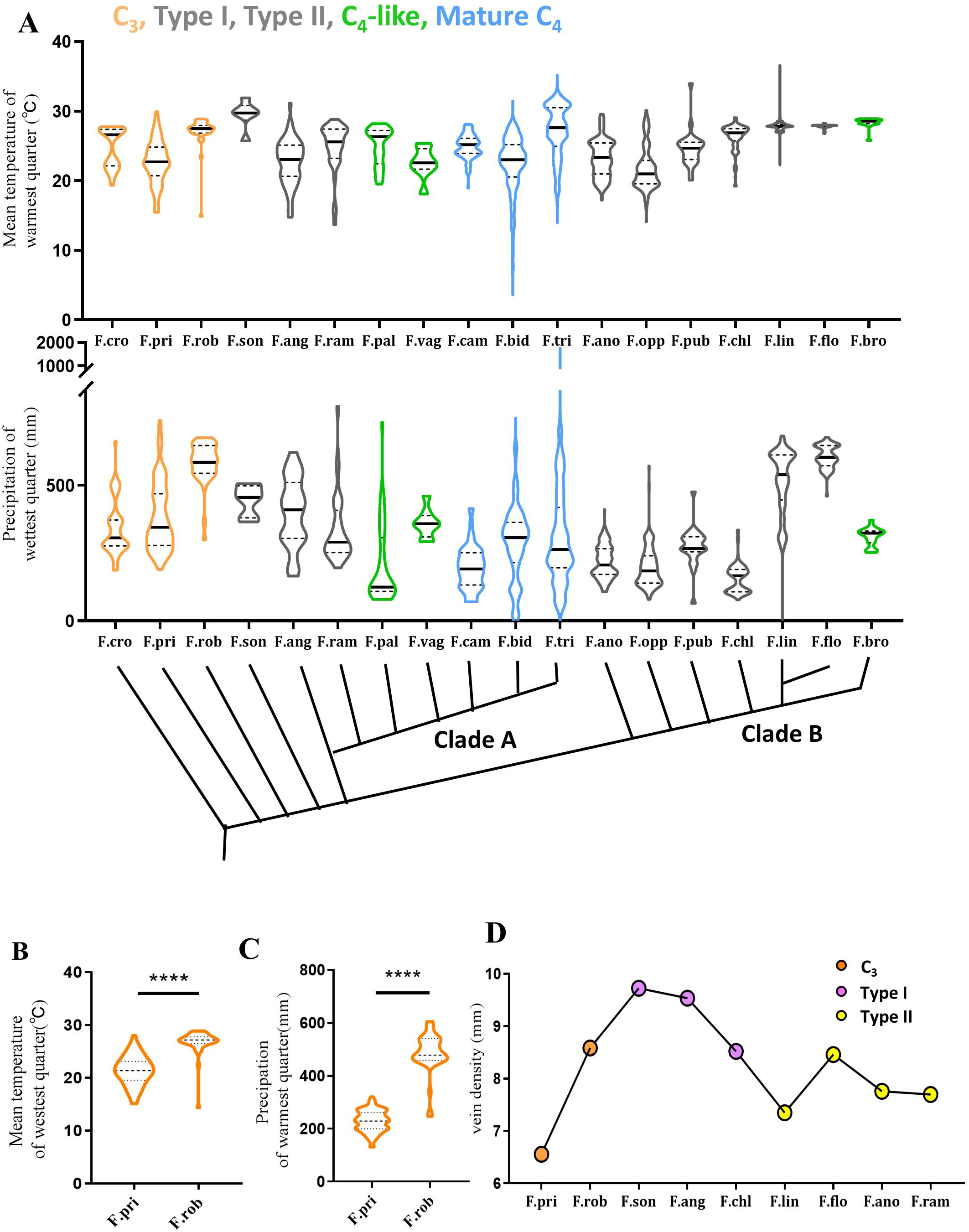
The environment conditions of the habitats for *Flaveria* species (A) The average temperatures of the warmest quarter for different *Flaveria* species (above panel). The precipitation of the wettest quarter for different *Flaveria* species (below panel). (B) The average temperatures of the wettest quarter for *F. rob* and *F. pri*. (C) The precipitation of wettest quarter for *F*.*rob* and *F*.*pri*. (D) The changes in vein density of C_3_ and intermediate species. Different colors indicate different photosynthetic types. Orange color represents C_3_, grey color represents intermediate species, green color represents C_4_-like, blue color represents C_4_ species. Black lines represent median values. These environmental data are from 30 years. *F*.*rob* (n=39), *F*.*cro* (n=48), *F. pri* (n=140), *F*.*son* (n=6), *F*.*ang* (n=50), *F*.*ano* (n=145), *F*.*opp* (n=135), *F*.*pub* (n=25), *F*.*ram* (n=65), *F*.*chl* (n=96), *F*.*flo* (n=56), *F*.*lin* (n=799), *F*.*cam* (n=95), *F*.*bro* (n=19), *F*.*bid* (n=525), *F*.*pal* (n=44), *F*.*tri* (n=1652), *F*.*vag* (n=8). For (A), the statistic test for the difference in the environmental conditions was performed with One-way ANOVA, and the difference between each group was performed by multiple Tukey test. For (B, C), the statistic test for the difference between *F. rob* and *F. pri* was performed with Student t-test (Unpaired, Two-tails).

**Fig 3.**
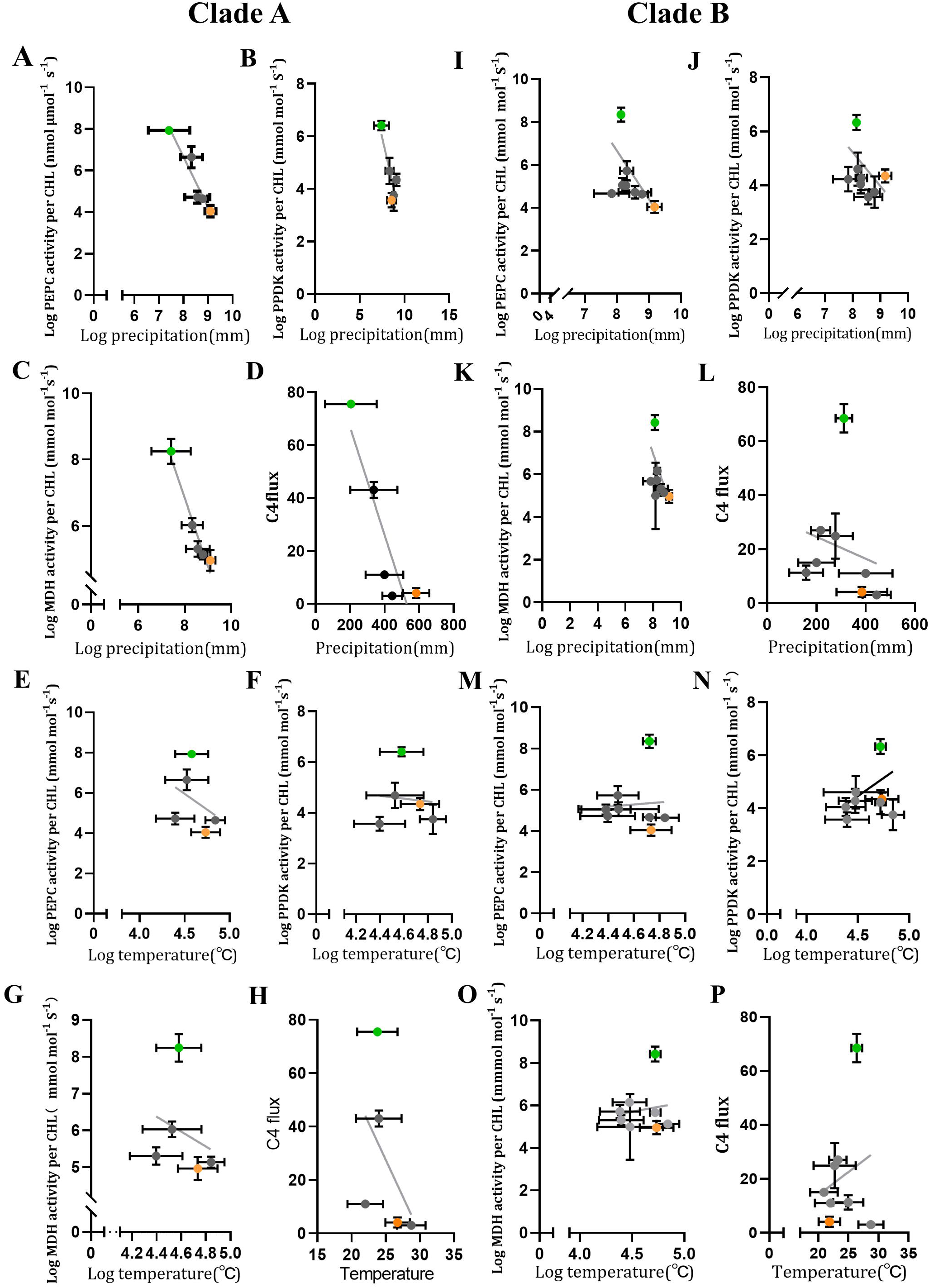
The association between precipitation and temperature with C_4_ enzyme activities or C_4_ flux Here the precipitation denotes the precipitation of the wettest quarter. The temperature denotes the average temperature of the wettest quarter. A-H, clade A. (A) The relationship between Log PEPC activity per CHL and Log precipitation. (B) The relationship between Log PPDK activity per CHL and Log precipitation. (C) The relationship between Log MDH activity per CHL and Log precipitation. (D) The relationship between C_4_ flux and precipitation. (E) The relationship between Log PEPC activity per CHL and Log temperature. (F) The relationship between Log PPDK activity per CHL and Log temperature. (G) The relationship between Log MDH activity per CHL and Log temperature. (H) The relationship between C_4_ flux and Log temperature. I-P, clade B. (I) The relationship between Log PEPC activity per CHL and log precipitation. (J) The relationship between Log PPDK activity per CHL and log precipitation. (K). The relationship between Log MDH activity per CHL and log precipitation. (L). The relationship between C_4_ flux and precipitation. (M) The relationship between log PEPC activity per CHL and log temperature. (N) The relationship between log PPDK activity per CHL and log temperature. (O) The relationship between log MDH activity per CHL and log temperature. (P) The relationship between C_4_ flux and temperature. The results of linear regression analysis are in supplementary file.

**Fig 4.**
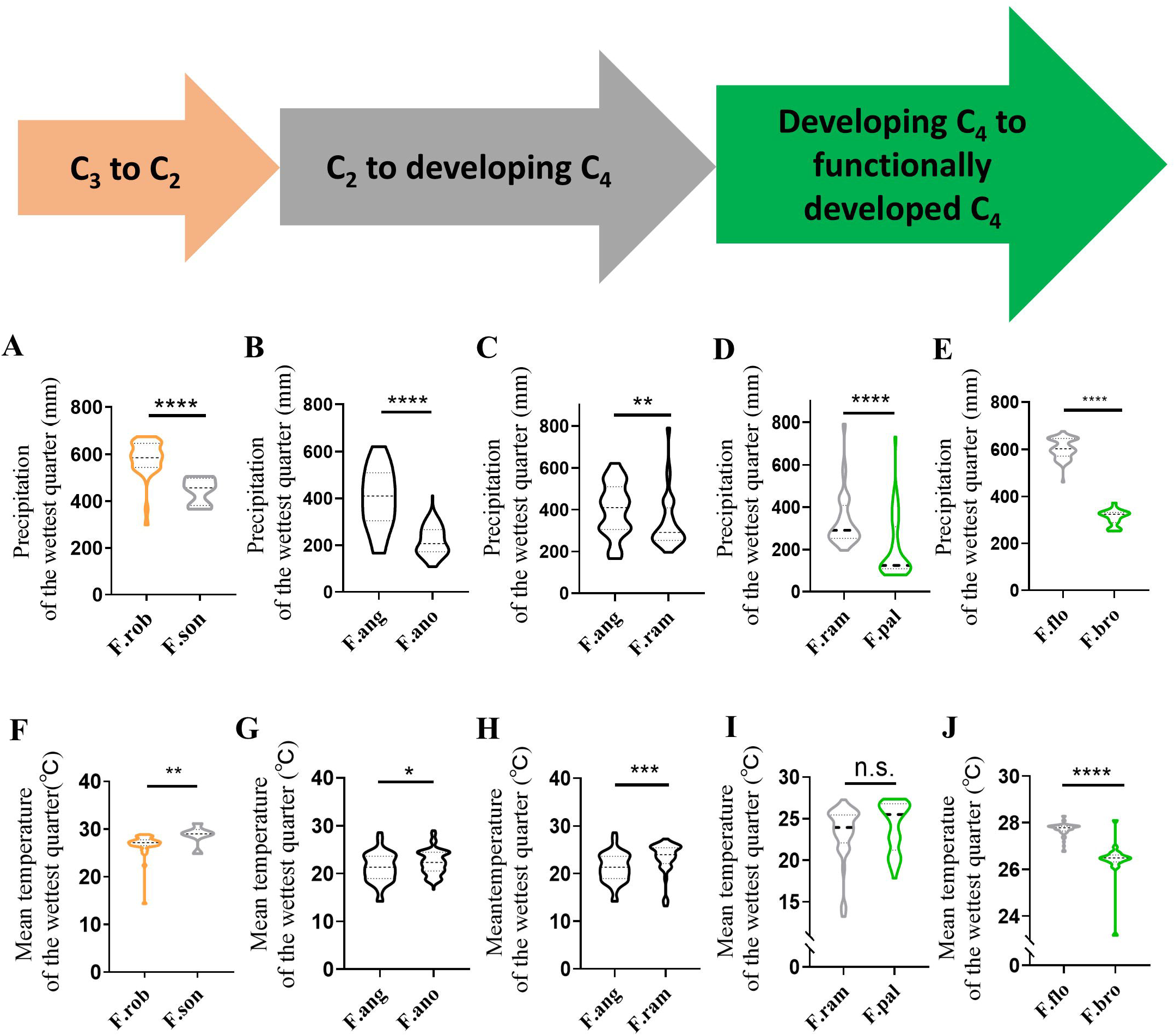
Precipitation and temperature during the transition from proto-Kranz to C_4_-like (A) Change in precipitation of wettest quarter during the transition from proto-Kranz (*F. rob*) to C_2_ (*F. son*). (B) Change in precipitation of wettest quarter during the transition from C_2_ (*F. son*) to developing C_4_ (*F. ano*) in clade B. (C) Change in precipitation of wettest quarter during the transition from proto-Kranz (*F*.*son*) to developing C_4_ (*F*.*ram*) in clade A. (D) Change in precipitation of wettest quarter during the transition from developing C_4_ (*F*.*ram*) to functionally developed C_4_ photosynthesis (*F*.*pal*) in clade A. (E) Change in precipitation of wettest quarter during the transition from developing C_4_ (*F*.*flo*)to functionally developed C_4_ photosynthesis (*F*.*bro*) in clade B. (F) Change in mean temperature of wettest quarter during the transition from proto-Kranz (*F*.*rob*) to C_2_ (*F*.*son*). (G) Change in mean temperature of wettest quarter during the transition from C_2_ (*F. son*) to developing C_4_ (*F. ano*) in clade B. (H) Change in mean temperature of wettest quarter during the transition from C_2_ (*F. son*) to developing C_4_ (*F. ram*) in clade A. (I) Change in mean temperature of wettest quarter during the transition from developing C_4_ (*F*.*ram*) to functionally developed C_4_ photosynthesis (*F. pal*) in clade A. (J) Change in mean temperature of the wettest quarter during the transition from developing C_4_ (*F. ano*) to functionally developed C_4_ photosynthesis (*F. bro*) in clade B. The transition from C_3_ to C_2_ and the emergence of C_4_ flux was promoted by increased temperature and decreased precipitation, indicated by the direction of arrow. That the arrow become broad indicates the enhancement of C_4_ flux. The statistic test for the difference between each group was performed with Student t-test (Unpaired, Two-tails).

For the precipitation of wettest quarter, precipitation of warmest quarter and precipitation of wettest month, along C_4_ evolutionary pathway, they significantly increased in the *F. rob* (C_3_) habitat, compared with those in *F. pri* habitat (Fig 2A, B, supplementary fig 3, supplementary file). There was an increased trend from preconditioning to proto-Kranz (Fig 2A, B, supplementary fig 3). Compared with *F. rob* (C_3_) habitat, precipitation significantly decreased in the *F. ang* (Type I) habitat (Fig 2A, supplementary fig 3, supplementary file). Precipitation decreased in *F. son* (Type I) habitat, and though the difference between *F. rob* and *F. son* is not statistically significant (Fig 2A, supplementary fig 3, supplementary file). Precipitation of warmest quarter was higher than precipitation of coldest quarter, and similar to precipitation of wettest quarter (Fig 2A, supplementary fig 2, supplementary figure 3, supplementary file). Expect *F. flo* (Type II) and *F. lin* (Type II), precipitation of the habitat of species with C_2_ photosynthesis decreased persistently along the path of C_4_ evolution, though difference for the habitats of some species is not significant (Fig 2A, supplementary fig 3, supplementary file). In the initialization of C_4_ evolution, the initial increase in precipitation and temperature (*F. pri* to *F. rob*) coincided with the initial appearance of the higher vein density (Fig 2A, B, C, supplementary fig 3, supplementary 4). Although the precipitation of the Type II species *F. flo* and *F. lin* habitat was high (Fig 2, supplementary fig 3, supplementary file), they distributed along coast and inhabited the saline land (Table 1), likely experiencing strong osmotic stress.

**Table 1.** The habitat of *Flaveria* species.

| Species<br>(photosynthetic type) | Growth habitat | Geographic<br>location |
| --- | --- | --- |
| *Flaveria pringlei*<br>(C<sub>3</sub>) | Rather arid and gypseous habitats, scattered localities, waste places, cultivated fields, roadsides | Southern Mexico |
| *Flaveria cronquistii*<br>(C<sub>3</sub>) | Rocky limestone (possibly gypseous) areas | Southern Mexico |
| *Flaveria robusta*<br>(C<sub>3</sub>) | Gypsum and slate, sandy clay, or open rocky slopes, weedy at roadsides | Western Mexico |
| *Flaveria sonorensis*<br>(Type I) | Close to warm mineral springs | North-western Mexico |
| *Flaveria angustifolia*<br>(Type I) | Sandy or loamy soils in open fields | Southern Mexico |
| *Flaveria pubescens*<br>(Type II) | Scattered localities, gypsum soils | Central Mexico |
| *Flaveria oppositifolia*<br>(Type I) | high gypsum plains or other areas with dilute gypsum, saline, or rarely loamy soil, usually in moist areas or in disturbed habitats at roadside | Central Mexico |
| *Flaveria linearis*<br>(Type II) | Near the coast and on offshore islands | Western Florida, USA |
| *Flaveria floridana*<br>(Type II) | Restricted to saline, white sand near the beaches of brackish marshes | Western Florida, USA |
| *Flaveria ramosissima*<br>(Type II) | Sandy-clay bottoms, along dry washes, fields and roadsides, possibly in gypseous soils, | Southern Mexico |
| *Flaveria anomala*<br>(Type II) | High gypsum plains, often in disturbed sites | Central Mexico |
| *Flaveria palmeri*<br>(C<sub>4</sub>-like) | Gypsum flats and knolls, or clay-loam soils probably with dilute gypsum, bolsons near lagunas, distributed areas near roads and fields | Northern Mexico |
| *Flaveria brownie*<br>(C<sub>4</sub>-like) | Saline, sandy, and marshy areas of the coastal flats and islands of the lower Texas Gulf Coast | North-eastern Mexico |
| *Flaveria vaginata*<br>(C<sub>4</sub>-like) | Arid and gypseous soils, scrub vegetation, cultivated fields | South-central Mexico |
| *Flaveria bidentis*<br>(C<sub>4</sub>) | A weed in moist places on waste ground, near rivers, streams, and barrancas, in fields, pastures, disturbed ground, near streets and sidewalks in towns, claygravel, or sand | Africa, Caribbean, South America, southern USA |
| *Flaveria trinervia*<br>(C<sub>4</sub>) | a variety of habitats, usually saline, gypseous, and disturbed, usually near ephemeral or permanent water sources | Africa, Caribbean, Central/South America, Mexico, USA |
| *Flaveria chloraefolia*<br>(Type I) | Permanent or ephemeral saline marshy soils, springs, streams, roadsides | North Mexico |

The information of habitat of *Flaveria* species was from (Powell 1978; Kocacinar et al. 2008)

### 3. The relationship between precipitation (or temperature) and C_4_ metabolism related traits during the transition from proto-Kranz to C_4_-like

For the clade A, the activities of C_4_ metabolism related enzymes PEPC, PPDK and MDH showed a negative correlation with precipitation, and there showed a linear relationship between log enzyme activity and log precipitation (Fig 3 A, B, C, supplementary file). The negative relationship between C_4_ flux and precipitation can be described as a linear function (Fig 3D). For the clade B, the relationship between log enzyme activity and log precipitation also can be described as a linear function (Fig 3 I, J, K, supplementary file). However, the linear relation between log enzyme activity and log precipitation in clade B is weaker than clade A (Fig 3I,J,K, supplementary file). A weak and negative relationship between C_4_ flux and precipitation was observed in clade B (Fig 3L). The linear relation between log enzyme activity and log temperature is weaker than that between log enzyme activity and log precipitation, regardless of clade A or clade B (Fig 3E, F, M, N, G, H, O, P, supplementary file). The linear relationship between C_4_ flux and temperature is weak in both clade A and clade B (Fig 3H, P).

### 4. The change in temperature and precipitation between the transition of stages during C_4_ evolution

The comparison of *F. rob* and *F. son* suggested that incipiently decreased precipitation along the evolutionary way towards C_4_ photosynthesis concurrently occurred with increased mean temperature (Fig 4A, F). Precipitation levels of the habitat of *F*.*ano* and *F*.*ram* were significantly lower than that of *F. ang*, and temperature values of the habitat of *F. ram* and *F. ano* were significantly higher than that of *F. ang* (Fig 4B, G, C, H). Precipitation of *F. pal* habitat was significantly lower than that of *F. ram*, and precipitation of *F. bro* habitat is significantly lower than that of *F. flo* (Fig 4D, E). However, temperature for the habitat of *F*.*pal* and *F. bro* showed the inconsistent change, temperature for the habitat of *F. pal* was similar to that of *F. ram*, and temperature of *F. bro* habitat was significantly lower than that of *F. flo* (Fig 4I, J). Furthermore, compared with their more ancient species, we observed that the decreased precipitation for the species habitat (*F. son, F*.*ram, F*.*flo, F*.*pal* and *F*.*bro*) concurrently appeared with the decreased stomatal density, decreased inhibition of O_2_ and CO_2_ compensation point, decreased maximum stomatal conductance (Supplementary Fig 5E, F, G, H). Although the precipitation of *F. flo* and *F. lin* were increased compared to other species with C_2_ photosynthesis (Fig 2A, supplementary fig3, supplementary fig4), they also had the decreased stomatal density, maximum stomatal conductance (Supplementary fig 5), which might be related to their salty habitats (Table1).

**Fig 5.**
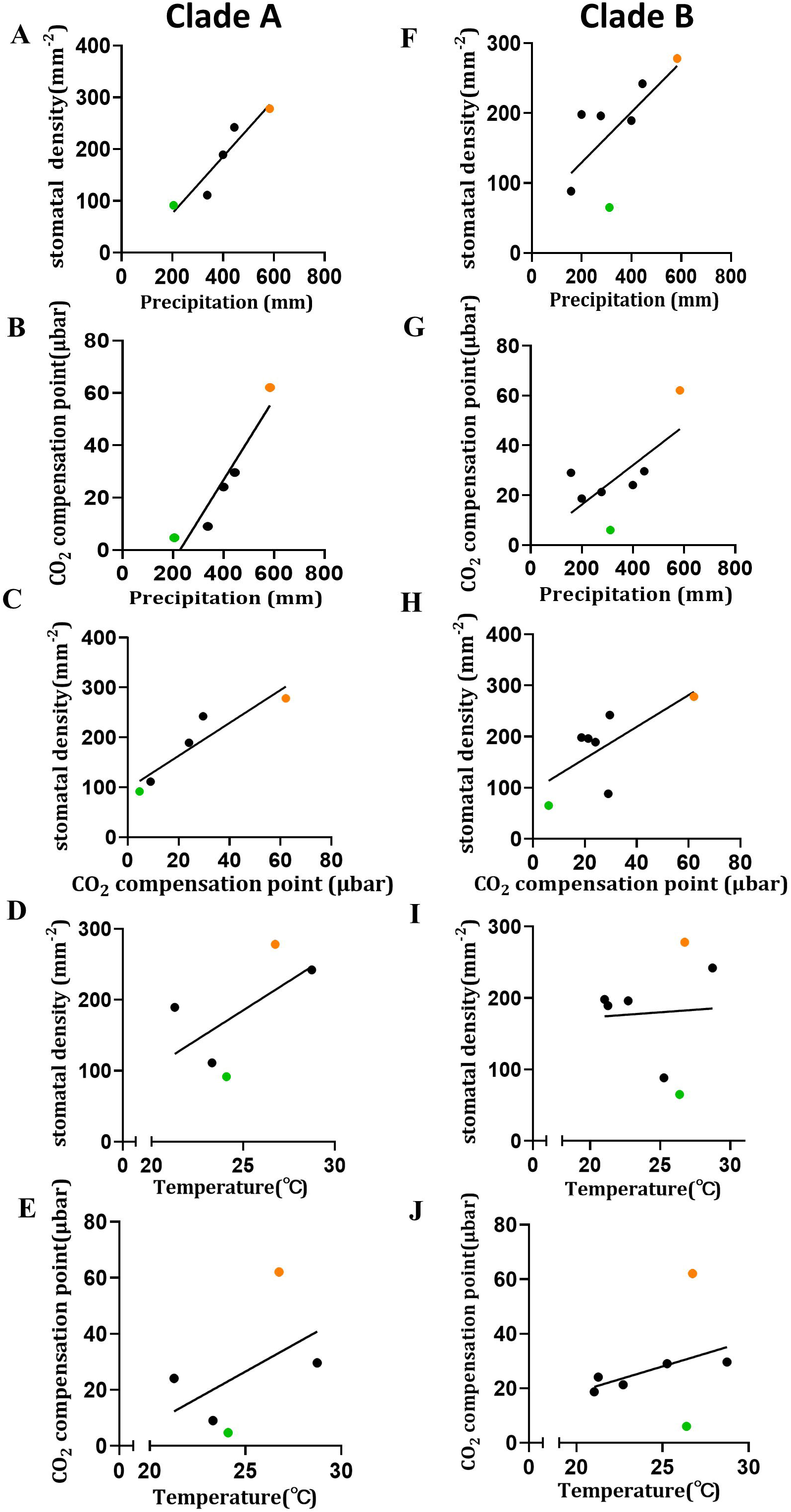
The association for stomatal density, CO_2_ compensation point, precipitation and temperature Here the precipitation denotes the precipitation of the wettest quarter. The temperature denotes the average temperature of the wettest quarter. A-E, clade A. (A) The relationship between stomatal density and precipitation. (B) The relationship between CO_2_ compensation point and precipitation. (C) The relation between stomatal density and CO_2_ compensation point. (D) The relationship between stomatal density and temperature. (E) The relationship between CO_2_ compensation point and temperature. F-J, clade B. (F) The relationship between stomatal density and precipitation. (G) The relationship between CO_2_ compensation point and precipitation. (H) The relationship between stomatal density and CO_2_ compensation point. (I) The relationship between stomatal density and temperature. (J) The relationship between CO_2_ compensation point and temperature. The data of stomatal density was from (Zhao et al. 2022), and the data of CO_2_ compensation point was from (Ku et al. 1991). The average of stomatal density and the average of CO_2_ compensation point were used in this study. The results of linear regression analysis are in supplementary file.

### 5. The relationship between precipitation, stomata and CO_2_ compensation point

Previous work found there was a difference in stomata density between species at different C_4_ evolutionary stages in the *Flaveria* genus (Zhao et al. 2022). Furthermore, there was a positive and linear relationship between stomata density, CO_2_ compensation point and precipitation during the transition between proto-Kranz and C_4_-like species in *Flaveria* clade A and clade B (Fig 5A, B, C, D, E, and F, Supplementary file). The linear relationship between stomata, CO_2_ compensation point and precipitation in *Flaveria* clade B were weaker than that in clade A (Fig 5A, B, C, F, G, H, Supplementary file). The relation between temperature and stomata density or CO_2_ compensation point was not apparent as the relationships between these two parameters and precipitation (Fig 5D, I, E, J, supplementary file).

## Discussion

### Evolutionary conditions along the path towards C_4_ photosynthesis

Heat and drought have long been thought to be associated with C_4_ evolution (Sage et al. 2018; Taylor et al. 2012). Exploring environmental conditions of the habitats for C_4_ and intermediate species could contribute to elucidate the drivers for C_4_ evolution (Sage et al. 2018; Sage et al. 2012; Edwards and Smith 2010). In this work, we examined precipitation and temperature in diverse *Flaveria* species during different periods (Fig 2A, supplementary fig 2, 3, 4). We noticed that the quantity and trend of wettest quarter mostly coincided with those of warmest quarter (Fig 2A, supplementary fig 3), mean temperature of wettest quarter was overall higher than mean temperature of driest quarter (Supplementary fig 3D, Supplementary fig 2E) and precipitation of warmest quarter was overall higher than precipitation of coldest quarter (Supplementary fig 3C, supplementary fig 2B). These results suggested that rain and heat were in the same season for the habitat of *Flaveria*. Sufficient water content was necessary to maintain high photosynthetic activity for plants, therefore this suggestion was in line with previous work that indicates *Flaveria* species perform high photosynthesis during summer with heat in Mexico (Powell 1978). Because there is no apparent change in Rubisco Sc/o (Kubien et al., 2008) and similar Ci/Ca between C_3_ and type I intermediate species (Adachi et al., 2023), the higher photosynthetic activity would indicate also a higher photorespiratory flux in C_3_ plants and initial intermediate species. Besides, under such conditions, photorespiration, long regarded as the driving factor of C_4_ evolution, might further be enhanced by heat (Sage et al. 2012; Sage et al. 2018). Furthermore, any evolutionary selective pressure on photosynthesis should act in the periods with active photosynthesis (Sage et al. 2018), i.e. the driest and coldest season should not be considered for the study on the evolutionary trajectory towards C_4_ photosynthesis (Supplementary fig 2). Therefore, the periods with rain and heat for *Flaveria* genus are valuable to explore the process of the driving of C_4_ evolution (Fig 2A, supplementary fig 3).

We found that the initial stage of C_4_ evolution was associated with high precipitation and temperature, and later stage was associated with decreased precipitation (Fig 2A, Fig 4, supplementary fig 3). Heat is considered to be a major factor that promotes the origin of C_4_ photosynthesis (Sage et al. 2018). Here, our results suggested that in addition to increased temperature, the increase in precipitation was also a vital factor to initiate the evolution towards C_4_ photosynthesis (Fig 2, Supplementary fig 3). It is possible that heat occurred with drought to enhance photorespiration simultaneously and then promoted C_4_ evolution (Sage et al. 2012). Here, data from this study indicated that this process indeed existed, and it was at the transition from C_3_ to C_2_ (*F. son*) and the transition from C_2_ to developing C_4_ photosynthesis (*F. ram, F. ano*) (Fig 4).

### The evolutionary mechanism during the initial phase of C_4_ evolution

The initial stage of C_4_ evolution is featured as having increased vein density and increased number of chloroplasts in bundle sheath (Sage et al. 2012). The increased vein density can help maintain sufficient water content in leaf and avoid the reduction of stomatal conductance, since the evaporation might be increased at this stage (Sage et al. 2012; Sage et al. 2018). In agreement with this, our data showed that, when at the stage of vein density increasing, there was no change in stomatal development, and temperature was indeed increased (Fig 2A, B, C, D, supplementary fig 5). The increased temperature could increase vapor pressure between leaf and air and hence enhance the evaporation (Sage et al. 2018). However, we found that water vaper pressure in atmosphere was increased in this stage and this could decrease water vapor deficit (Supplementary fig 6B). No change in the stomatal density and maximum stomatal conductance indicated that water shortage in leaf did not occur (Supplementary fig 5), since long-term water shortage could alter stomatal development (Franks et al. 2009; Taylor et al. 2012). Importantly, our data also showed that precipitation was increased at this stage, which matched with the increased vein density, and this ensured the sufficient supply of water to the leaf (Fig 2A, B, C). Increased precipitation and keeping stomatal opening could not only maintain intercellular CO_2_ concentration, but also could help alleviate the increase of leaf temperature. These changes all benefit the acquirement of carbon. Therefore, the initial stage of C_4_ evolution could be promoted by high temperature and precipitation (Fig 2).

**Fig 6.**
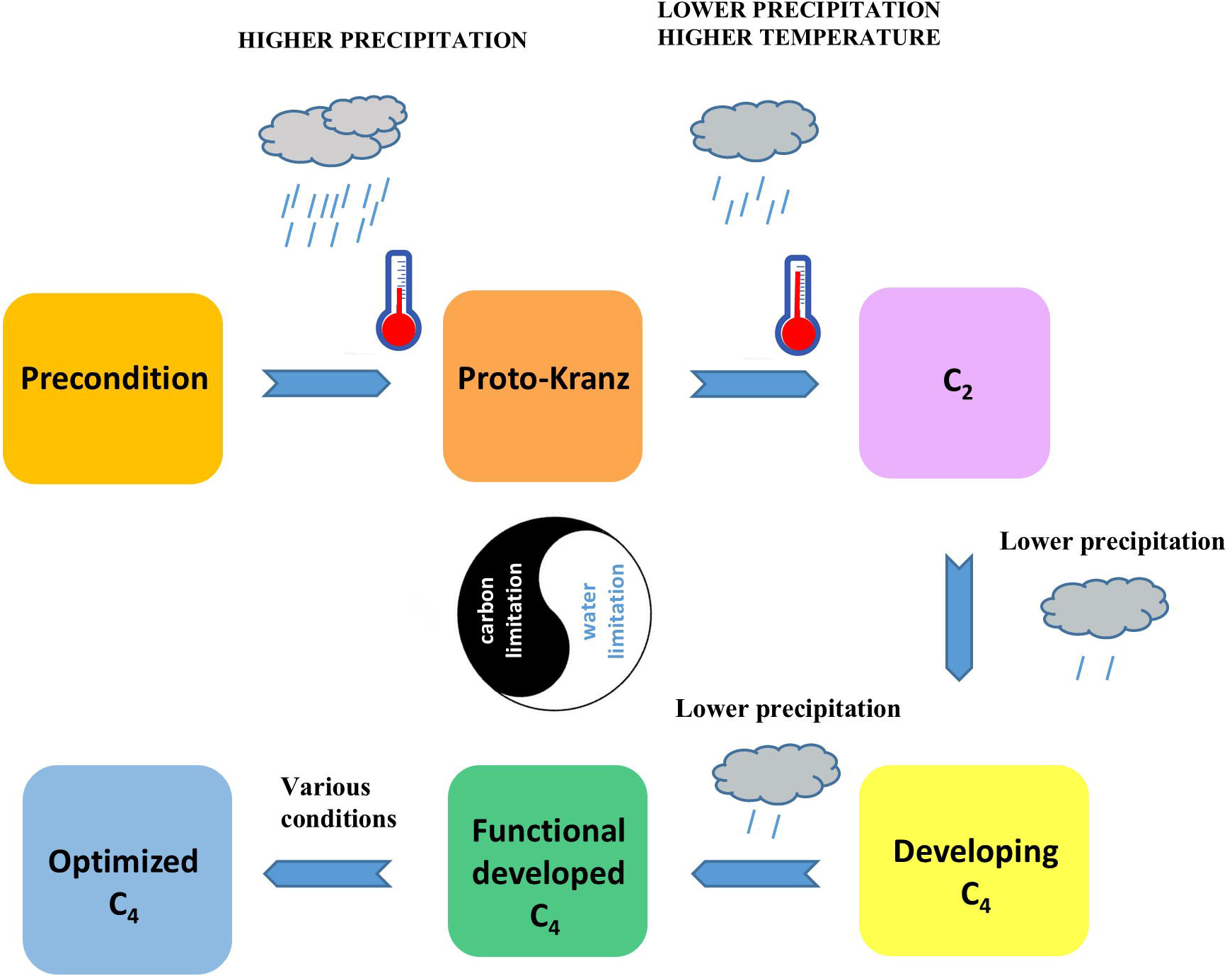
Schematic diagram for the driving of C_4_ evolution. The schematic diagram for the conditions driving the evolutionary process for C_4_ photosynthesis. The raindrop number denotes the precipitation. The thermometer denotes the temperature. The arrows denote the direction of C_4_ evolution. The cycle composed by white and black denotes the primary driving element that promote the corresponding stage of C_4_ evolution. The primary driving reason in C_3_ and proto-Kranz was carbon limitation, and the primary driving reason in the latter stage was water limitation. The black become narrow indicating the carbon limitation was increasing gradually, and the white become narrow indicating the water limitation was increasing gradually.

The increase in vein density could decrease volume of mesophyll in a leaf (Sedelnikova et al. 2018; Mckown and dengler 2007), which inevitably decreases the photosynthetic capacity of the leaf. In theory, this should create a strong evolutionary pressure to select for the increased photosynthetic capacity of BSC to offset the loss of photosynthetic capacity. In consistent with this, the relative number and size of chloroplast in bundle sheath cell of *F. rob* is slightly increased compared to those of *F*.*pri* (Sage et al. 2013). Besides, the photosynthetic apparatus located into BSC from MC could benefit the refixation of CO_2_, since BSC locates to the inner of leaf and photosynthetic apparatus in BSC locates to the inner cell periphery (Sage et al. 2013). Indeed, the Proportion in half of cell, coverage of inner half of cell area and compensation CO_2_ point at high light intensity are also slightly increased (Sage et al. 2013). The light intensity of *F. rob* habitat was higher than that of *F. pri* (Supplementary fig 5A); the size of bundle sheath in *F. rob* decreases compared with *F. pri* (Sage et al. 2013; Mckown and dengler 2007); the size of mesophyll cell decreases compared with *F. pri* (Mckown and dengler 2007); and the thickness of leaf become less (Mckown and dengler 2007); all these enhanced the level of light that bundle sheath experiences, consisted with the notion that selection pressure for bundle sheath with higher photosynthetic capacity might exist at this stage.

### Drought as a more important factor than heat for driving C_4_ evolution

Some studies have proposed that drought is a primary driving force for C_4_ evolution (Sage 2004; Zhou et al. 2018; Sage et al. 2018; Schluter and Weber 2016; Edwards and Smith 2010), and other studies suggest that heat is the critical environment factor driving C_4_ evolution (Sage et al. 2018; Sage et al. 2012). In this study, we proposed that drought played a more important role than heat during C_4_ evolution (Fig 3). From proto-Kranz to C_4_-like, precipitation declined step by step, however, there was no clear trend of change in temperature (Fig 2). Regardless of clade A or Clade B, we found a negative association between core C_4_ enzyme activity and precipitation (Fig 3, supplementary file) and positive association between CO_2_ compensation point and precipitation (Fig 5, supplementary file). In particular, for clade A, there was a strong negative relationship between precipitation and the activities of core C_4_ enzymes (Fig3, supplementary file). However, the relationship wasn’t apparent for temperature (Fig 3, supplementary file). These results indicated that progressing C_2_ photosynthesis and developing C_4_ photosynthesis mainly associated to decreasing precipitation (Fig 2A, B, C, D, E, F, Fig 3). Therefore, drought rather than heat, was suggested to be the major driving force for the evolution from proto-Kranz to C_4_-like. Flaveria genus has long been thought as the chief model in C_4_ evolutionary study and it represents C_4_ evolution of all evolutionary route to a large extent. Therefore, drought might dominate this transitional in all C_4_ evolutionary route. C_4_ evolution repeat independently above 60 times (Sage et al. 2012). It is possible that heat dominates this transitional process in the evolution of other C_4_ lineages. In future, the drivers for this transitional process in other C_4_ evolutionary lineages still need to be checked.

### Different evolutionary drivers at the transition of the stages in C_4_ evolution

The environmental drivers for the evolution towards C_4_ photosynthesis have been intensively studied. For example, decreased concentration of atmosphere CO_2_ has been thought as a driving force of C_4_ evolution (Christin et al. 2008; Concentrations and Paleogene). Drought, salt and heat were also thought to be conditions that promote C_4_ evolution (Sage et al. 2018; Sage 2004; Bromham and Bennett 2014) (Edwards and Still 2008; Edwards and Smith 2010; Pearcy and Ehleringer 2006; Sage et al. 2012). These studies assume that entire C_4_ evolutionary process is promoted by a single ecological condition. However, here, our results suggested that environmental conditions promoted C_4_ evolution in an orderly and stepwise manner (Fig 6), and different selection pressure drove the transition of certain stages during the evolutionary way towards C_4_ photosynthesis (Fig 6).

Besides the initialization of C_4_ evolution, the evolution towards C_2_ photosynthesis is also considered to be a crucial step during C_4_ evolution (Sage et al. 2012; Schluter and Weber 2016). Previous studies have reported that *Flaveria* species with C_2_ photosynthesis can represent a transition between C_3_ and C_4_ photosynthesis during the evolution (Heckmann et al. 2013; Mallmann et al. 2014; Schluter and Weber 2016; Sage et al. 2012), and *F. ram* and *F. ano* have enhanced C_4_ cycle flux, C_4_ cycle related enzyme activity and gene expression levels (Adachi et al. 2023; Mallmann et al. 2014; Vogan and Sage 2011; R.K. Monson 1986). According to the phylogenetic tree, *F. ang* is closer to *F*.*ram* as compared with *F*.*son* (Fig 2A), therefore, among all *Flaveria* species, *F*.*son* most likely represents the emergence of C_2_ photosynthesis in *Flaveria*. Similarly, *F. ram* and *F. ano* most likely represent the emergence of developing C_4_ photosynthesis in clade A and clade B, respectively, and *F. pal* and *F. bro* most likely represent the emergence of a fully functional C_4_ cycle for clade A and clade B, respectively (Fig 2A). Therefore, the increased temperature and decreased precipitation simultaneously promoted the emergence of C_2_ photosynthesis (Fig 4A, F). Increasing temperature and decreasing precipitation could enhance photorespiration and thus promote the formation of photorespiratory CO_2_ pump which improves the fitness of these intermediate plants. The photorespiratory CO_2_ pump largely decreases the photorespiration (Sage et al. 2018; Sage et al. 2012; Schluter and Weber 2016; Ku et al. 1991), which is indicated by the large decrease in CO_2_ compensation point and O_2_ inhibition (Sage et al. 2013; Ku et al. 1991). Study proposes photorespiration promotes C_4_ evolution by decreasing stomatal conductance under drought (Sage et al. 2012). As the support, we provided the evidence that there was a positive relationship between stomatal density, CO_2_ compensation point and precipitation (Fig 5). These also suggested that the concurrent evolution for decreased stomata density and the emergence of C_2_ and C_4_ photosynthesis during C_4_ evolution (Fig 5). The enhancement of C_2_ photosynthesis has long been considered to cause the imbalance of nitrogen between mesophyll cell and bundle sheath cell, which induces C_4_ cycle (Mallmann et al. 2014). In case of the evolutionary process from C_2_ photosynthesis to developing C_4_ photosynthesis, precipitation decreased and temperature increased (Fig 4B, C, G, H), and this evolutionary process could be driven. The formation of functionally developed C_4_ photosynthesis was driven by inconsistent conditions for Clade A and Clade B (Fig 4I, J), and precipitation have played a dominated role in this step. There was no significant difference and a decrease in temperature of the habitats (Figure 4 I, J), which suggested that the temperature was not a deciding factor controlling this transition.

In summary, this study analyzed the data of ecology, physiology and biochemistry for different species of *Flaveria* genus to reveal the process that the evolution from C_3_ towards C_4_ photosynthesis is driven, and results indicated plants suffered carbon limitation in the early phase of C_4_ evolution and the primary driving force for this phase is carbon limitation (Fig 6). Later, plants suffered water limitation leading to decease in acquirement of carbon, therefore, the primary driving force for these phases of C_4_ evolution is water limitation (Fig 6). For a long time, most of studies proposed that environmental condition causing the carbon restriction through enhancing photorespiration (drought, heat, salty and low concentration of atmosphere CO_2_) drove the evolution towards C_4_ photosynthesis(Christin et al. 2008; Edwards and Smith 2010; Schluter and Weber 2016; Sage 2004; Sage et al. 2012; Sage et al. 2018). Our study proposed these studies were not subtlety-oriented and suggested that certain steps of the evolution from C_3_ to C_4_ photosynthesis were driven by different environmental factors which sequentially promoted C_4_ evolution, in contrast to the current notion that the entire process of C_4_ evolution is driven by same environmental factor (Fig 6). The initial evolution was promoted by high precipitation, while later decreased precipitation positively dominated the transition among C_3_, C_2_ and C_4_ photosynthesis (Fig 6). More important, the increase in vein density could represent a crucial characteristics for the starting point of the evolution from C_3_ towards C_4_ photosynthesis (Christin et al. 2012; Mckown and dengler 2007; Sage et al. 2012), therefore, increased humidity and temperature together may be the most important condition that facilitated the initial evolution towards C_4_. Therefore, the origin of C_4_ evolution might be in the habitat with raised humidity and temperature, rather than drought, heat, salt and low concentration of atmosphere CO_2_ (Bromham and Bennett 2014; Christin et al. 2008; Sage et al. 2018). Our study could benefit the prediction of the occurrence of C_4_ evolution. C_4_ photosynthesis is crucial for the ecology and environment. Therefore, this study may help us handle the climate and environment changes currently and in the future. Additionally, the understanding provided in this study for the driving of C_4_ evolution might benefit to identify the regulatory innovation that facilitated C_4_ evolution, which might help us to perform C_4_ engineering.

## Methods

### 1. Obtain the ecological data of *Flaveria* species

First, the geographical distribution of *Flaveria* was acquired from Global Biodiversity Information Facility (GBIF) database using the rgibf software package. Using ArcMAP software, we acquired from the Worldclim database the elevation data of the respective locations and 19 Bioclimatic factors, including Mean Diurnal Range, annual mean Temperature, Isothermality, Temperature Seasonality, Max Temperature of Warmest Month, Temperature Annual Range, Mean Temperature of Wettest Quarter, Mean Temperature of Warmest Quarter, Annual Precipitation, Precipitation of Wettest Month, Precipitation of Warmest Quarter, Precipitation of Wettest Quarter, Precipitation of Coldest Quarter, Min Temperature of Coldest Month, Precipitation of Driest Quarter, Mean Temperature of Coldest Quarter, Precipitation of Driest Month, Precipitation Seasonality (Coefficient of Variation), Precipitation of Coldest Quarter based on the geographical distribution information of the *Flaveria*. Mean Diurnal Range, Isothermality, Temperature Seasonality, Temperature Annual Range, Precipitation Seasonality (Coefficient of Variation) were not used in this study. Specifically, we downloaded the data from the worldclim at first. Secondly, we prepared the file for the information of sampling location. Thirdly, we imported the layers file of climate factor. Fourthly, we imported the information of sampling location. Fifth, generating the new layers. Lastly, we exported the file that including sample and the data of geographical factors. All bioclimatic variables were averaged from 1970 to 2000. All data from the worldclim database have a maximum spatial resolution of 30 arc-seconds. The water vaper pressure and solar radiation were also extracted from the World clim database(Fick and Hijmans 2017), based on the geographical distribution information of *Flaveria*. The water vaper pressure and solar radiation in June, July and August were extracted. The water vaper pressure and solar radiation were also from 1970-2000.

### 2. Obtain the biochemical, anatomical and physiological data of *Flaveria* species

The biochemical, anatomical and physiological data for *Flaveria* was acquired from previous research. The data of vein density and distance between vein was from (Mckown and dengler 2007). The data of vein density and distance between vein was digitized by Origin 2018. Specifically, first, importing the figure with the picture format; Second, compiling the coordinate axis with the Image digitizing tool in Origin 2018; Third, grabbing and obtaining the data point with the Image digitizing tool in Origin 2018; Fourth, exporting the data. The data of CO_2_ compensation point and O_2_ inhibition was from (Ku et al. 1991), and the means of the data of CO_2_ compensation point and O_2_ inhibition were used in this study. The data related to stomata was from our work (Zhao et al. 2022), and the means were used in this study. The data of C_4_ metabolic enzyme activity was from (Supplementary table S7) in (Adachi et al. 2023). The growth habitat was acquired from (Powell 1978; Kocacinar et al. 2008).

### 3. Map the *Flaveria* species

We mapped and marked the locations for the habitats of different *Flaveria* species to a world map using the software google earth Pro (version 7.3) based on the Latitude and longitude from Global Biodiversity Information Facility (GBIF). Coordinates were plotted as placemarks, with customized icons to distinguish between different species to facilitate the visualization and analysis of distribution patterns and geographical clusters. Google Earth Pro, version 7.3. (2020). [Software]. Google LLC. Available at: https://www.google.com/earth/.

### 4. Statistical analysis, regression analysis and Graph plotting

The graph was plotted and the regression analysis was performed by using GraphPad prism 8. The statistic test was performed by GraphPad prism 8, WPS office or Microsoft office. The statistic test for the difference in precipitation or temperature between all *Flaveria* species was performed by one-way ANOVA, and the difference between each group was conducted using Tukey’s multiple test (Figure 2A, supplementary Figure 2,3,4, supplementary file). The rest of other statistic test (Figure 2B, C, Figure 4) was conducted by Student’s t test, and the detail in the statistic test was described in each figure legends. The linear regression in Figure 3 was conducted by using the log _2_ value of the precipitation of wettest quarter and log _2_ value of precipitation of C_4_ metabolic enzyme activities, and linear regression between them were performed with GraphPad prism 8 (Supplementary file). Specifically, conducting the option of analyze; Then, selecting linear regression; Exporting the result. The parameters in GraphPad prism 8 for the linear regression were the defaults. The results of the linear regression were in supplementary file. The phylogenetic tree for *Flaveria* was drawn by integrating the studies in (Lyu et al. 2015; Mckown 2005).

## Figure legends

Supplementary fig 1. The C_4_ flux for *Flaveria*. The data of C_4_ flux was from table 1 in (Vogan and Sage 2011)

Supplementary fig 2. Temperature and precipitation of habitat for *Flaveria* species in driest and coldest period. (A) Precipitation of the driest quarter. (B) Precipitation of the coldest quarter. (C) Precipitation of the driest month. (D) Mean temperature of the driest quarter. (E) Mean temperature of coldest quarter. (F) Min temperature of coldest month. Orange, C_3_; Purple, Type I; Yellow, Type II; Green, C_4_-like; Blue, C_4_. Black lines indicate median. The plants abbreviate genus written. *rob* (n=39), *cro* (n=48), *pri* (n=140), *son* (n=6), *ang* (n=50), *ano* (n=145), *opp* (n=135), *pub* (n=25), *ram* (n=65), *chl* (n=96), *flo* (n=56), *lin* (n=799), *cam* (n=95), *bro* (n=19), *bid* (n=525), *pal* (n=44), *tri* (n=1652), *vag* (n=8).

Supplementary fig 3. Temperature and precipitation of habitat for *Flaveria* species in wettest and warmest period. (A) Max temperature of warmest month. (B) Precipitation of westest month. (C) Precipitation of warmest quarter. (D) Mean temperature of wettest quarter. Orange indicates C_3_, purple indicates Type I, yellow indicates Type II, green indicates C_4_-like, blue indicates C_4_. Black lines indicate median. The plants abbreviate genus written. *rob* (n=39), *cro* (n=48), *pri* (n=140), *son* (n=6), *ang* (n=50), *ano* (n=145), *opp* (n=135), *pub* (n=25), *ram* (n=65), *chl* (n=96), *flo* (n=56), *lin* (n=799), *cam* (n=95), *bro* (n=19), *bid* (n=525), *pal* (n=44), *tri* (n=1652), *vag* (n=8).

Supplementary fig 4. Annual temperature and precipitation of habitats for *Flaveria* species (A) Annual mean temperature. (B) Annual precipitation. Orange, C_3_; Purple, Type I; Yellow, Type II; Green, C_4_-like; Blue, C_4_. Black lines indicate median. The plants abbreviate genus written. *rob* (n=39), *cro* (n=48), *pri* (n=140), *son* (n=6), *ang* (n=50), *ano* (n=145), *opp* (n=135), *pub* (n=25), *ram* (n=65), *chl* (n=96), *flo* (n=56), *lin* (n=799), *cam* (n=95), *bro* (n=19), *bid* (n=525), *pal* (n=44), *tri* (n=1652), *vag* (n=8).

Supplementary fig 5. The trend in features for the evolution towards C_4_ photosynthesis. (A) Precipitation of wettest quarter. (B) Mean temperature of wettest quarter. (C) Vein density. (D) Distance between vein. (E) O_2_ inhibition. (F) CO_2_ compensation point. (G) Maximum stomatal conductance (G_smax_) (H) Stomatal density (SD). The plants abbreviate genus written.

Supplementary fig 6. The difference in solar radiation and water vaper pressure between *F. pri* and *F. rob*. A, The difference in solar radiation between *F*.*pri* and *F*.*rob*. B, The difference in water vaper pressure between *F*.*pri* and *F*.*rob. F. rob* (n=39). *F. pri* (n=140). Table 1 The habitat for *Flaveria* species

Supplementary file 1: The statistics test and regression analysis. The plants abbreviate genus written.

## Acknowledgements

Thanks for the research group, and the postdoc support in shanghai. National Research and Development Program of Ministry of Science and Technology of China (2020YFA0907600)

## Author contribution

YYZ and ZGZ designed the study. YXZ and YYZ extracted the climate data and the latitude and longitude data, and YYZ analyzed these climate data. XQY and YYZ mapped and marked the coordinates of *Flaveria* genus location and do the visualization. YYZ extracted the physiological data from the previous works. YYZ analyzed all these data, and did the statistic test and regression analysis. YYZ plotted the graphs. YYZ and XGZ wrote the paper.

