## Supplementary figures and images for "The process that the evolution from C_3_ towards C_4_ photosynthesis is driven"

### Supplementary figure 1

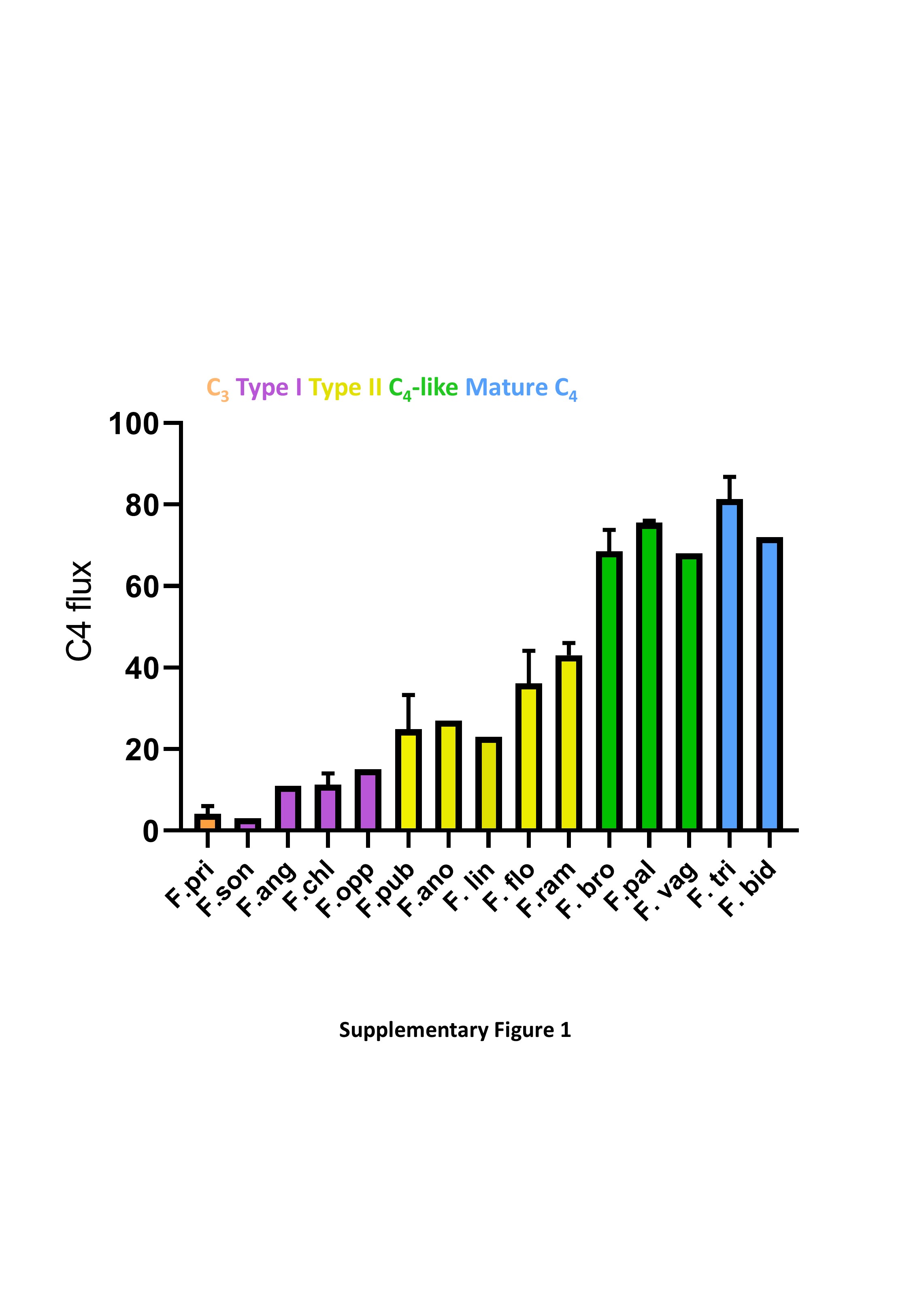

### Supplementary figure 2

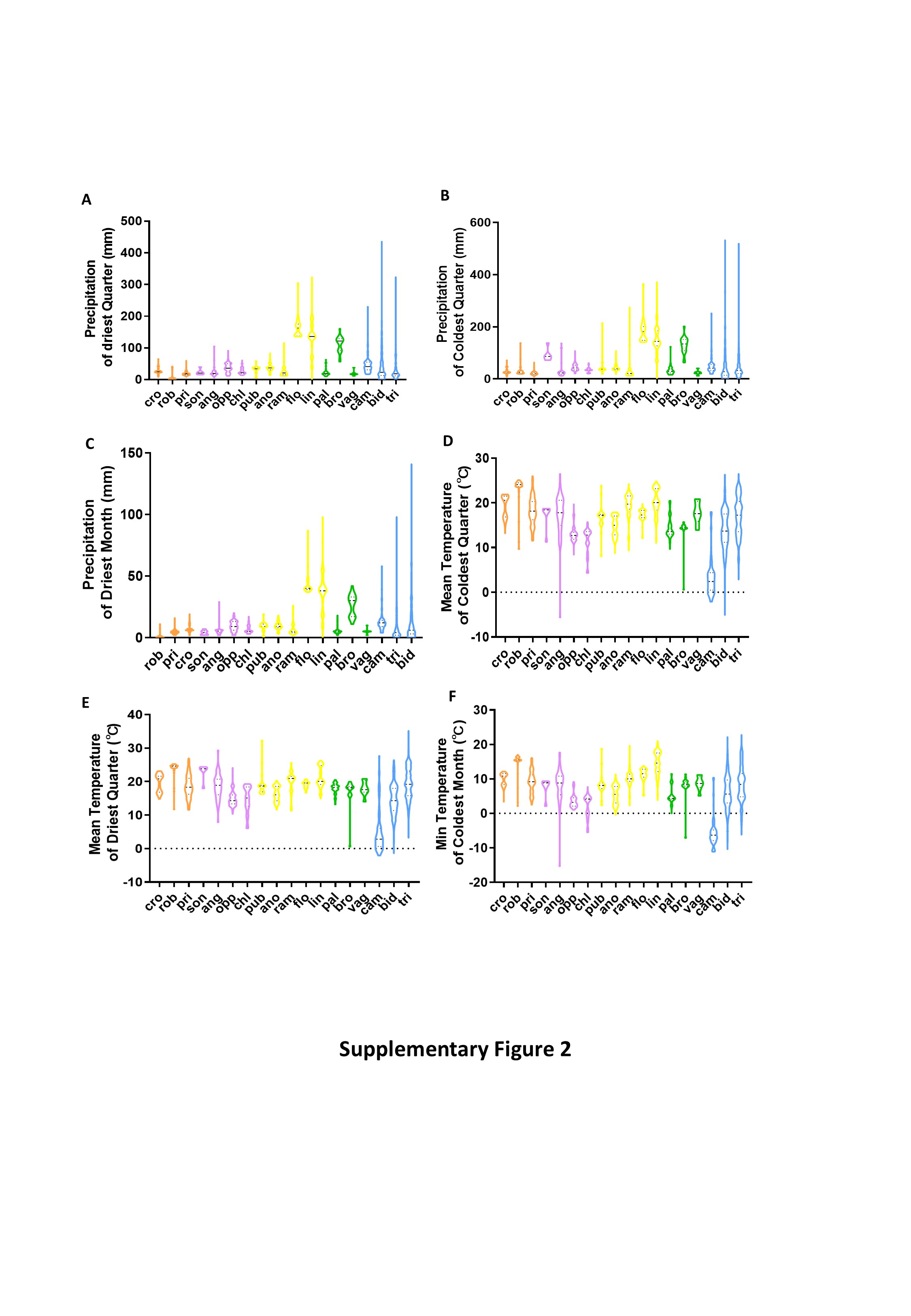

### Supplementary figure 3

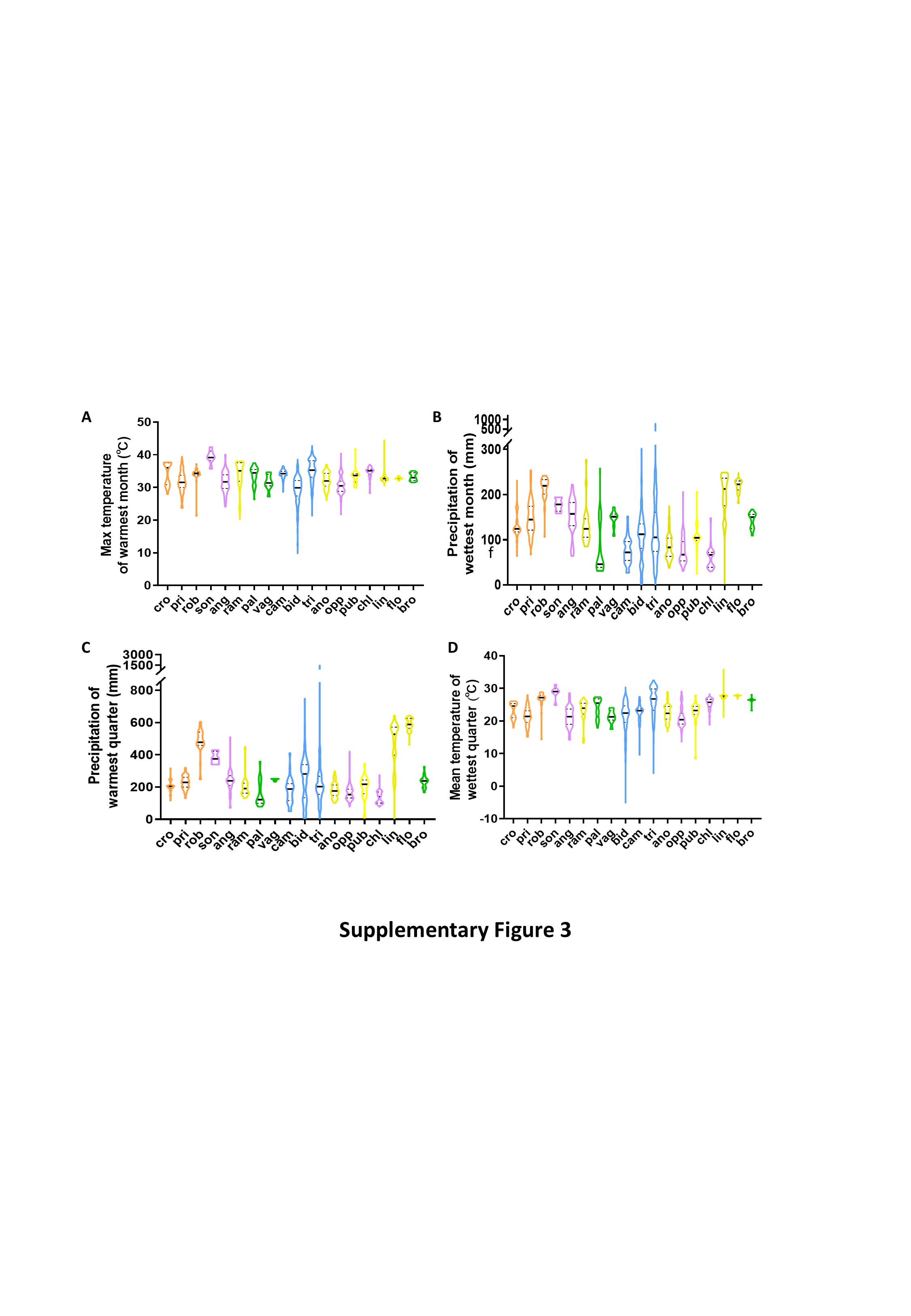

### Supplementary figure 4

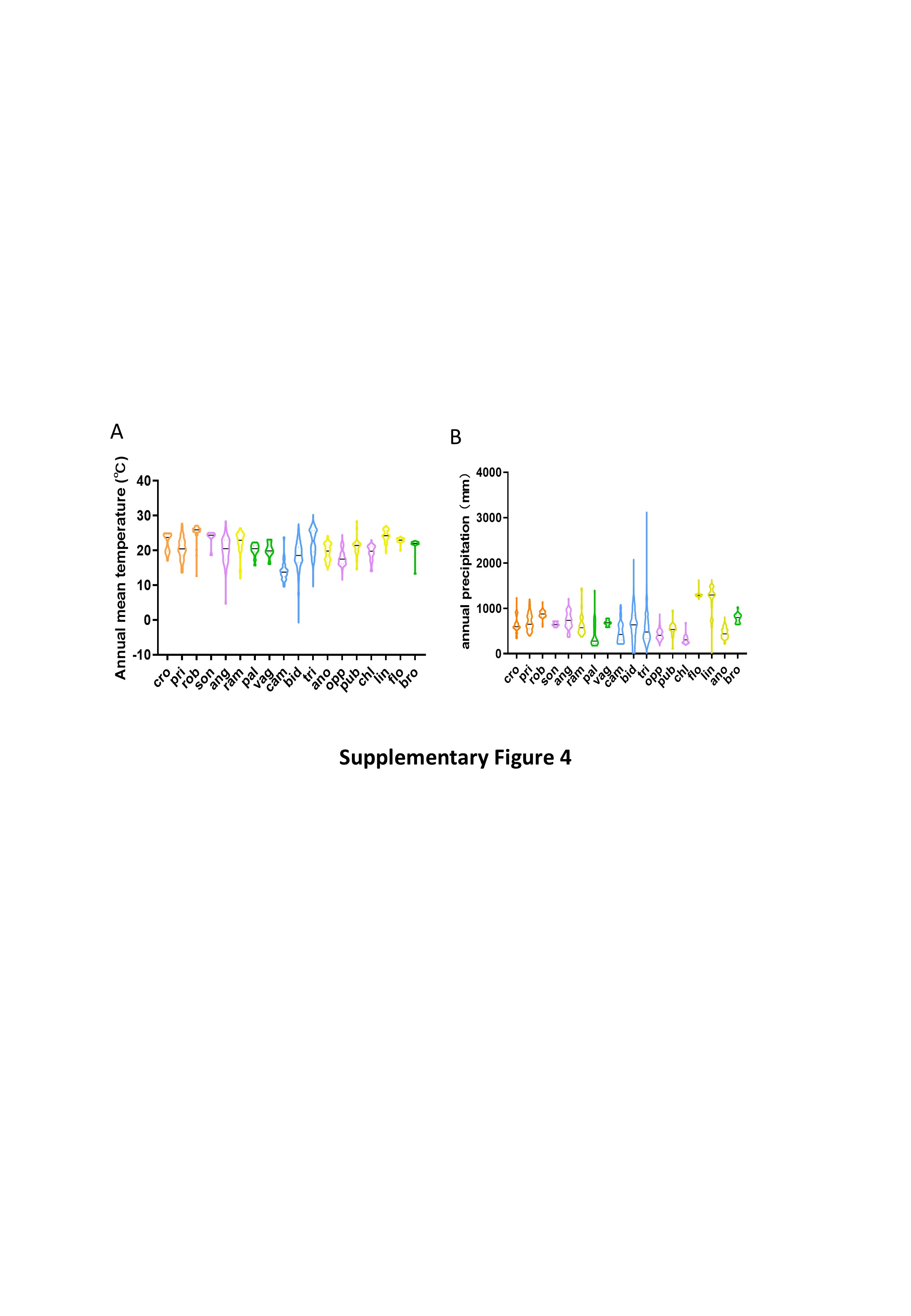

### Supplementary figure 5

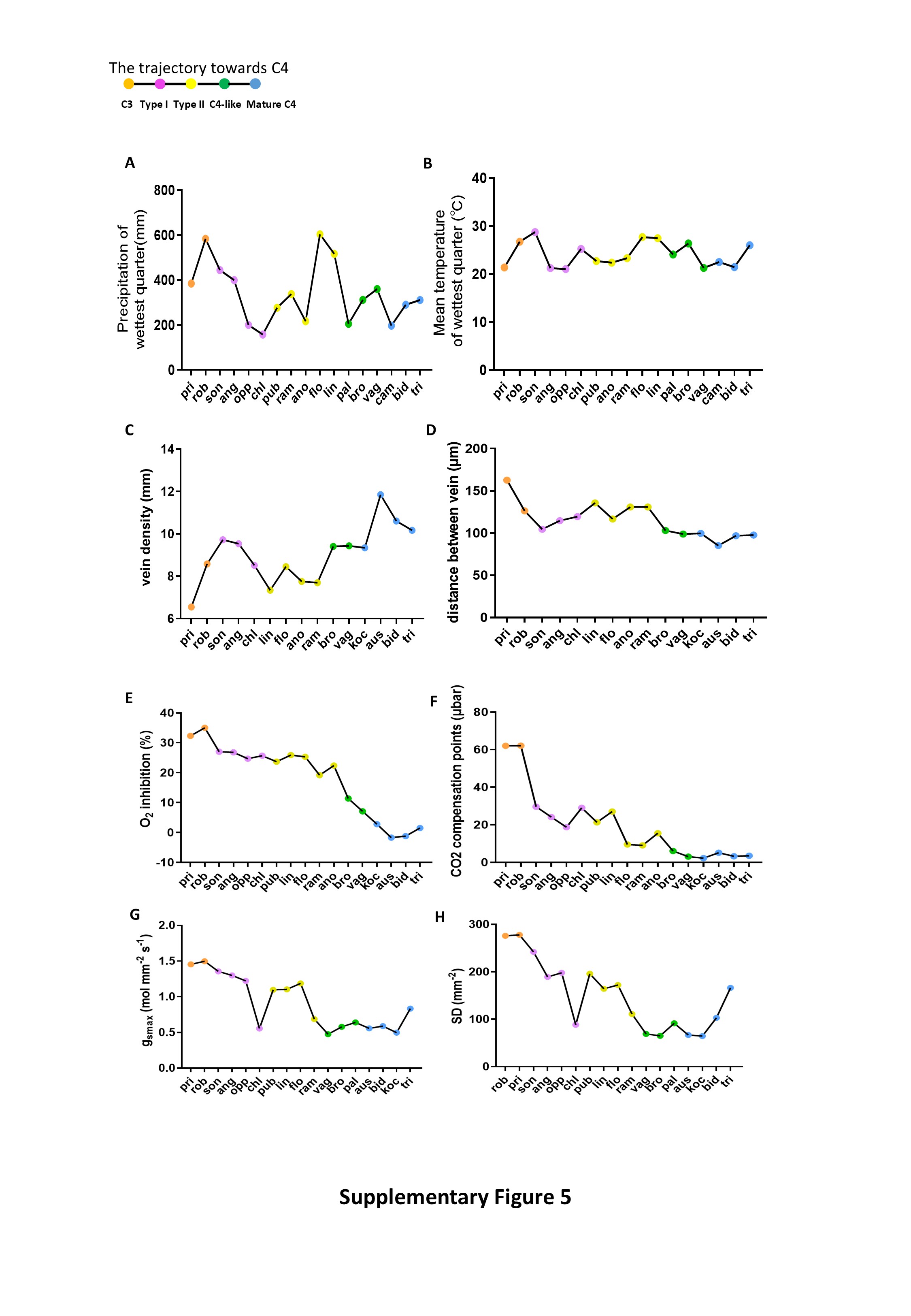

### Supplementary figure 6

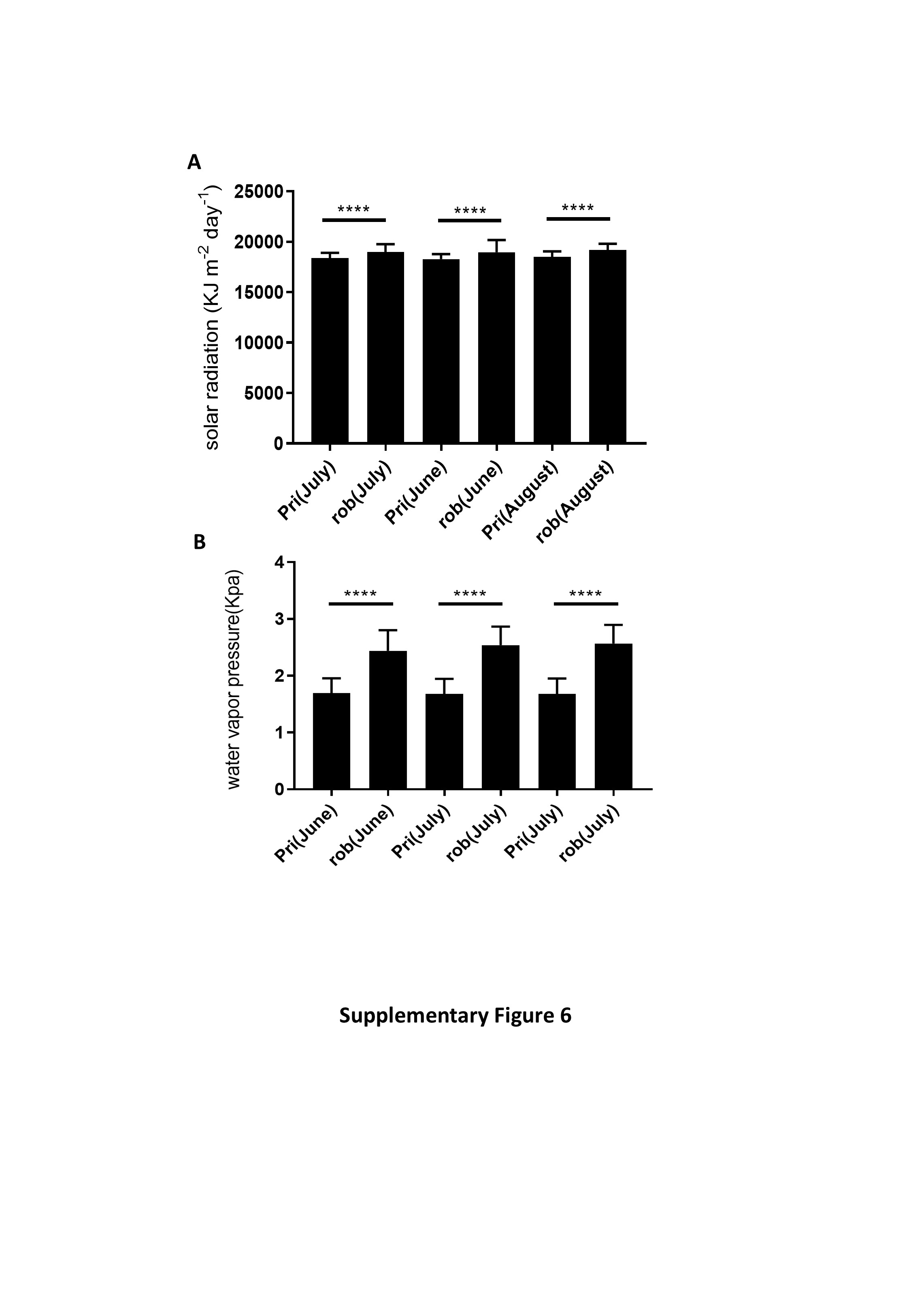
